# Predicting microRNA targeting efficacy in Drosophila

**DOI:** 10.1101/198689

**Authors:** Vikram Agarwal, Alexander O. Subtelny, Prathapan Thiru, Igor Ulitsky, David P. Bartel

**Author notes:** **Corresponding author**. Whitehead Institute for Biomedical Research, 9 Cambridge Center, Cambridge, MA 02142, USA.

## Abstract

Important for understanding the regulatory roles of miRNAs is the ability to predict the mRNA targets most responsive to each miRNA. Here, we acquired datasets needed for the quantitative study of microRNA targeting in Drosophila. Analyses of these data expanded the types of sites known to be effective in flies, expanded the mRNA regions with detectable targeting to include 5′ UTRs, and identified features of site context that correlate with targeting efficacy. Updated evolutionary analyses evaluated the probability of conserved targeting for each predicted site and indicated that more than a third of the Drosophila genes are preferentially conserved targets of miRNAs. Based on these results, a quantitative model was developed to predict targeting efficacy in insects. This model performed better than existing models and will drive the next version of TargetScanFly (v7.0; targetscan.org), thereby providing a valuable resource for placing miRNAs into gene-regulatory networks of this important experimental organism.

## INTRODUCTION

MicroRNAs (miRNAs) are ∼22-nt regulatory RNAs that originate from hairpin precursors (Bartel 2004). In Drosophila, they associate primarily with the Argonaute1 (dmAgo1) protein to form a silencing complex (Forstemann et al. 2007; Tomari et al. 2007), within which the miRNA functions as a sequence-specific guide that recognizes target mRNAs through pairing to complementary sites primarily within the 3′ untranslated regions (3′ UTRs) (Lai 2002; Brennecke et al. 2005; Bartel 2009).

The miRNA pathway found in flies is ancestral to animals (Grimson et al. 2007), with dozens of miRNA genes conserved broadly in bilaterian species (Ruby et al. 2007; Lu et al. 2008; Mohammed et al. 2013; Fromm et al. 2015). Small-RNA sequencing has identified hundreds of miRNAs that are encoded in fly genomes (Lagos-Quintana et al. 2001; Aravin et al. 2003; Ruby et al. 2007; Berezikov et al. 2011; Kozomara and Griffiths-Jones 2014; Fromm et al. 2015), which in aggregate appear to target thousands of mRNAs (Brennecke et al. 2005; Grun et al. 2005; Rehwinkel et al. 2006; Kheradpour et al. 2007; Ruby et al. 2007; Schnall-Levin et al. 2010; Berezikov et al. 2011; Jan et al. 2011). Studies of miRNAs in *Drosophila melanogaster* have helped define components of the miRNA processing pathway, developmental roles of miRNAs, and evolutionarily conserved mechanisms of miRNA action (Smibert and Lai 2010).

Drosophila miRNAs are expressed in complex spatiotemporal patterns throughout development (Sempere et al. 2003; Aboobaker et al. 2005) and play a wide diversity of roles. Examples include functions for the bantam miRNA in the regulation of cell proliferation (Brennecke et al. 2003), miR-iab4/iab8 in body patterning (Bender 2008; Stark et al. 2008; Tyler et al. 2008) and behavior (Picao-Osorio et al. 2015), miR-14 in insulin production and metabolism (Varghese et al. 2010), miR-34 in aging and neurodegeneration (Liu et al. 2012), and miR-277 in branched-chain amino acid catabolism (Esslinger et al. 2013). Indeed, a large-scale survey of miRNA knockouts in the flies reports abnormal knockout phenotypes for more than 80% of the miRNA genes tested (Chen et al. 2014).

Crucial for understanding the molecular basis of these phenotypes is the search for, and characterization of, miRNA targets. Analyses of reporter assays and site conservation indicate that the canonical site types identified in mammals, which include perfect Watson–Crick pairing to the miRNA seed (miRNA nucleotides 2–7) (Lewis et al. 2005), also function in flies (Brennecke et al. 2005; Grun et al. 2005; Lai et al. 2005; Kheradpour et al. 2007; Ruby et al. 2007; Stark et al. 2007; Schnall-Levin et al. 2010; Jan et al. 2011). However, knowledge of miRNA targeting in flies has lagged behind that of mammals, primarily due to the lack of high-throughput datasets examining the responses of mRNAs to the perturbation of miRNAs. In mammals, such datasets have been very useful for both measuring the relative efficacy of different site types and identifying additional features that influence site efficacy, such as those related to the context of the site within the mRNA, thereby enabling the development of quantitative models of site efficacy (Bartel 2009). Although as in mammals, much of miRNA targeting in flies is known to be seed-based, the relative importance of site types and context features might differ between mammals and flies, calling into question the utility for flies of quantitative models developed using mammalian data. For instance, fly 3′ UTRs are shorter and have a higher AU-content than those of mammals, which would presumably affect the utility of context features such as distance from a 3′-UTR end or local AU content, which are known to be predictive of site efficacy in mammals (Grimson et al. 2007). Although some attempt to model the effect of target-site accessibility on miRNA-mediated repression has been applied to Drosophila as well as mammals (Kertesz et al. 2007), the relatively poor performance of this model when tested in mammalian systems suggests that in fly it would have also benefited from the use of large datasets for training and validation (Baek et al. 2008).

Despite the lack of high-throughput repression data, many algorithms have been developed to predict and rank miRNA targets in *Drosophila*. Most, including EMBL predictions (Brennecke et al. 2005; Stark et al. 2005), EIMMo (Gaidatzis et al. 2007), MinoTar (also available as TargetScanFly ORF) (Schnall-Levin et al. 2010), miRanda-MicroCosm (Griffiths-Jones et al. 2008), PicTar (Grun et al. 2005; Anders et al. 2012), and TargetScanFly (Ruby et al. 2007), use a mix of pairing and evolutionary criteria, with pairing sometimes evaluated using predicted thermodynamic stability. Others, including PITA (Kertesz et al. 2007), RNA22 (Miranda et al. 2006), and RNAhybrid (Rehmsmeier et al. 2004), utilize purely thermodynamic information. Others, such as DIANA-microT-CDS (Reczko et al. 2012), mirSVR (Betel et al. 2010), and TargetSpy (Sturm et al. 2010), were trained on mammalian data using machine-learning strategies and then used to generate predictions for flies.

As with most algorithms applied in mammals, some of those applied in flies predict many non-canonical target sites that have one or more mismatches or wobbles to the miRNA seed. However, others, including DIANA-microT-CDS, EIMMo, MinoTar, RNAhybrid, and TargetScanFly, require perfect seed pairing in an effort to enhance the specificity of detecting functional targets, although it is unclear to what degree this comes at the price of reduced sensitivity. Whereas most algorithms limit predictions to sites in 3′ UTR’s, DIANA-microT-CDS and MinoTar also include predictions with sites in coding regions, which seem to have an even greater signal for preferential conservation in flies than they do in mammals (Lewis et al. 2005; Schnall-Levin et al. 2010).

Here, we used RNA-seq to monitor the effects of introducing specific miRNAs into Drosophila cells. Analyses of these data, together with updated analyses of site conservation in flies and other insects, provided new and quantitative insights into the types of target sites that function in flies, the scope of targeting in flies, and features of site context that influence site efficacy. With these insights, we generated a quantitative model that improves the rankings of target predictions for the fly miRNAs, which will be available at TargetScanFly, v7.0 (http://www.targetscan.org). We also release an accompanying suite of computational tools to help others reproduce our figures and apply our analyses to future datasets (TargetScanTools; https://github.com/vagarwal87/TargetScanTools).

## RESULTS & DISCUSSION

### Canonical miRNA target sites function primarily in *Drosophila* 3′ UTRs

To acquire datasets suitable for quantitative analysis of miRNA targeting in fly cells, we monitored the changes in mRNA levels after co-transfecting S2 cells with one of six different miRNA duplexes and a GFP-encoding plasmid. The six transfected miRNAs (miR-1, miR-4, miR-92a, miR-124, miR-263a, and miR-994) were chosen because they (or related miRNAs in the same seed family) were not endogenously expressed in S2 cells (Ruby et al. 2007), and they had diverse starting-nucleotide identities, a range of GC content within their seeds, and a moderate-to-high range of predicted target-site abundances. After enriching for transfected, GFP-positive cells by FACS, mRNA-seq was performed, and mRNA fold changes were calculated for each miRNA transfection condition relative to a mock transfection, in which the GFP plasmid was transfected without any miRNA duplex (Table S1). We then normalized the data to reduce batch effects (Figure S1) and began investigating the features of miRNA target sites that correlate with mRNA repression in Drosophila cells.

In mammals, the presence of an A opposite the first nucleotide of a miRNA is preferentially conserved and correlates with enhanced repression, regardless of the identity of the first nucleotide of the miRNA—observations explained by a pocket within human Argonaute2 (hsAGO2) that preferentially binds this A (Lewis et al. 2005; Grimson et al. 2007; Baek et al. 2008; Schirle et al. 2015). In flies, an A at this position of the target site is also associated with enhanced conservation compared to otherwise identical sites missing this A (Jan et al. 2011), whereas in nematodes conservation and efficacy of a sites with perfect pairing to miRNA nucleotides 2–8 followed by a U (8mer-U1 sites) resembles that of 8mer-A1 sites (Clark et al. 2010; Zisoulis et al. 2010; Jan et al. 2011). We therefore examined the influence of the nucleotide at target position 1 in flies, considering the data from all miRNA transfections pooled together. Of the mRNAs possessing a single match to miRNA nucleotides 2–8 in their 3′ UTR, those with an A opposite miRNA position 1 (i.e., those with the 8mer-A1 site) tended to be more repressed than those with each of the other three possibilities opposite miRNA position 1 (8mer-C1, 8mer-G1, and 8mer-U1, respectively), with the identity of the other three possibilities having little influence on repression (Figure 1A). As expected based on the observation that the first position of the guide RNA is buried within Argonaute and unavailable for pairing (Ma et al. 2005; Parker et al. 2005; Schirle et al. 2015), this observation generally held when considering each miRNA transfection independently, regardless of whether the identity of the first nucleotide of the miRNA was a U (Figure S2). Thus *Drosophila* exhibits a preference for A at target position 1 resembling that of mammals, implying that this target nucleotide is recognized by a pocket within dmAgo1 resembling that of hsAGO2. With respect to nomenclature, these results further supported consideration of the 8mer-A1 site as the canonical 8mer site of Drosophila, as was done originally in mammals (Lewis et al. 2005).

**Figure 1.**
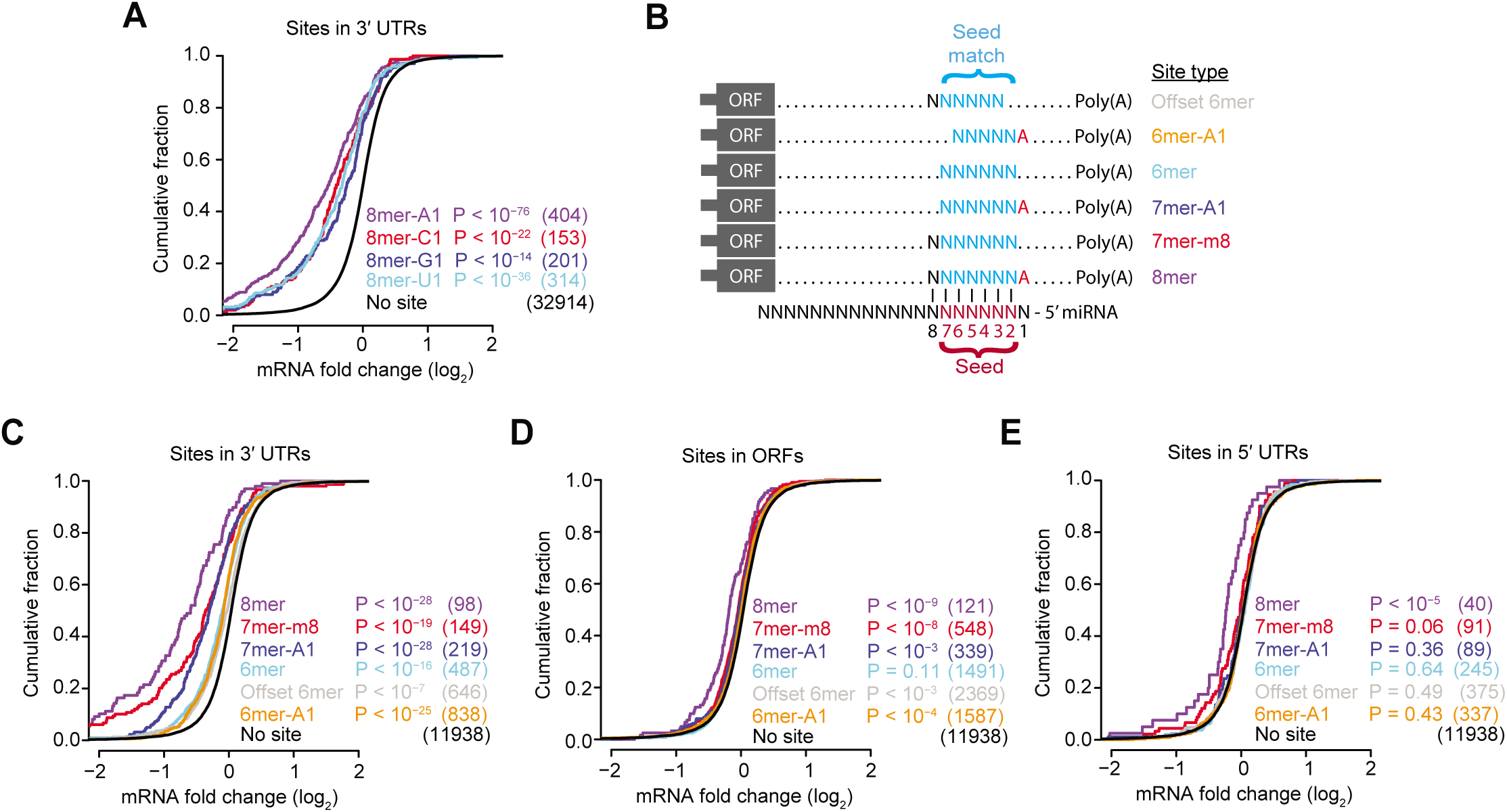
Drosophila miRNAs mediate mRNA repression through the targeting of canonical site types, preferentially in 3′ UTRs. **(A)** The increased efficacy in Drosophila of sites with an A across from miRNA position 1. Shown is the response of mRNAs to the transfection of a miRNA (either miR-1, miR-4, miR-92a, miR-124, miR-263a, or miR-994). Data were pooled across these six independent experiments. Plotted are cumulative distributions of mRNA fold changes observed upon miRNA transfection for mRNAs that contained a single site of the indicated type to the transfected miRNA. The site types compared are 8mers that perfectly match miRNA positions 2–7 and have the specified nucleotide (A, C, G, or U) across from position 1 of the miRNA. Also plotted for comparison is the cumulative distribution of mRNA fold changes for mRNAs that did not contain a canonical 7-or 8-nt site to the transfected RNA in their 3′ UTR (no site). Similarity of site-containing distributions to the no-site distribution was tested with the one-sided Kolmogorov–Smirnov (K–S) test (*P* values). Shown in parentheses are the numbers of mRNAs analyzed in each category. **(B)** The six canonical site types for which a signal for repression was detected after transfecting a miRNA into Drosophila cells. **(C–E)** The efficacy of the canonical site types observed in Drosophila 3′ UTRs **(C)**, ORFs **(D)**, and 5′ UTRs **(E)**. These panels are as in (A), but compare fold-change distributions for mRNAs possessing a single canonical site in the indicated region to those with no canonical sites in the entirety of the mRNA. See also Figure S1 and Figure S2.

Analogous analyses of mRNA fold-change values in mammalian systems have demonstrated the function and relative efficacy of 8mer, 7mer-m8, 7mer-A1, 6mer, and offset 6mer sites (Grimson et al. 2007; Friedman et al. 2009). Accordingly, we examined the function of these site types in *Drosophila*, again pooling the data and focusing on mRNAs with a single site to the cognate miRNA. We also considered a sixth site type, the 6mer-A1 site, which has implied function in nematodes (Jan et al. 2011) and completes the set of all possible 8-, 7-and 6-nt perfect matches to the 8- nt seed region, which we refer to as the canonical site types (Figure 1B). When located in the context of 3′ UTRs, each canonical site type tended to mediate repression, with the magnitude of repression following the hierarchy of 8mer > 7mer-m8 > 7mer-A1 > 6mer ∼ offset 6mer ∼ 6mer-A1 (Figure 1C). This hierarchy resembled that of mammals, except in mammals the efficacy of the different 6-nt sites is much more distinct, with 6mer > offset 6mer > 6mer-A1, and with the 6merA1 difficult to distinguish from background (Grimson et al. 2007; Friedman et al. 2009).

We also examined the efficacy of canonical sites in mRNA regions outside of the 3′ UTR. Some repression was observed for mRNAs with a site in their ORF (and no canonical site elsewhere in the mRNA), most convincingly for 8mer sites, although the efficacy of these sites was much less than that observed in 3′ UTRs (Figure 1D). These observations are consistent with those in mammals (Grimson et al. 2007; Gu et al. 2009; Schnall-Levin et al. 2011). In contrast to observations in mammals, however, repression was also observed for mRNAs with an 8mer site in their 5′ UTR (Figure 1E). Taking these findings together, we conclude that miRNA targeting in flies resembles that of mammals, except the efficacy of the three 6-nt canonical sites is more uniform in flies and repression of endogenous mRNAs is more readily detected in fly 5′ UTRs.

### Widespread conservation of canonical miRNA target sites in *Drosophila* UTRs

A previous evolutionary analysis of mammalian miRNA target sites provided a framework for estimating the likelihood that predicted miRNA target sites are conserved across species, while controlling for factors such as differential species relatedness, differential background conservation in untranslated regions, and differential rates of dinucleotide substitutions (Friedman et al. 2009). Although this approach has also been applied to *Drosophila* genomes (Jan et al. 2011), we improved and extended it by: i) updating conserved miRNA family classifications and 3′-UTR annotations, ii) using an expanded evolutionary tree that incorporated additional insect species, iii) extending analyses to *Drosophila* 5′ UTRs, iv) using a modified evolutionary analysis pipeline (Agarwal et al. 2015), and v) comparing our evolutionary results to our functional data. Towards this end, we compiled miRNA annotations from multiple studies (Ruby et al. 2007; Mohammed et al. 2013; Kozomara and Griffiths-Jones 2014; Fromm et al. 2015) and classified 91 miRNA families as broadly conserved among Drosophila species, 29 of which have been conserved since the last bilaterian ancestor (Table S2). We also extracted multiple sequence alignments corresponding to annotated *D. melanogaster* 5′ UTRs and 3′ UTRs, assigning each UTR to one of five bins based on its background UTR conservation rates (Jan et al. 2011). For each bin, we computed phylogenetic trees with a fixed species tree topology that encompassed 27 insect species, allowing for variable branch lengths to capture slower or faster substitution rates among the UTRs of the bin (Figure 2A). These trees were then used to assign a branch-length score (Kheradpour et al. 2007) to each motif occurrence in *D. melanogaster* UTRs, which quantified the extent of conservation of that occurrence while controlling for the background conservation rate of its overall UTR context (Friedman et al. 2009). For example, a motif occurrence detected among all Sophophora species in the 3′ UTR alignment would be assigned a branch-length score of 4.50, 2.53, or 1.69, depending upon whether the corresponding 3′ UTR in which it resided was in the first, third, or fifth conservation bin, respectively (Figure 2A).

**Figure 2.**
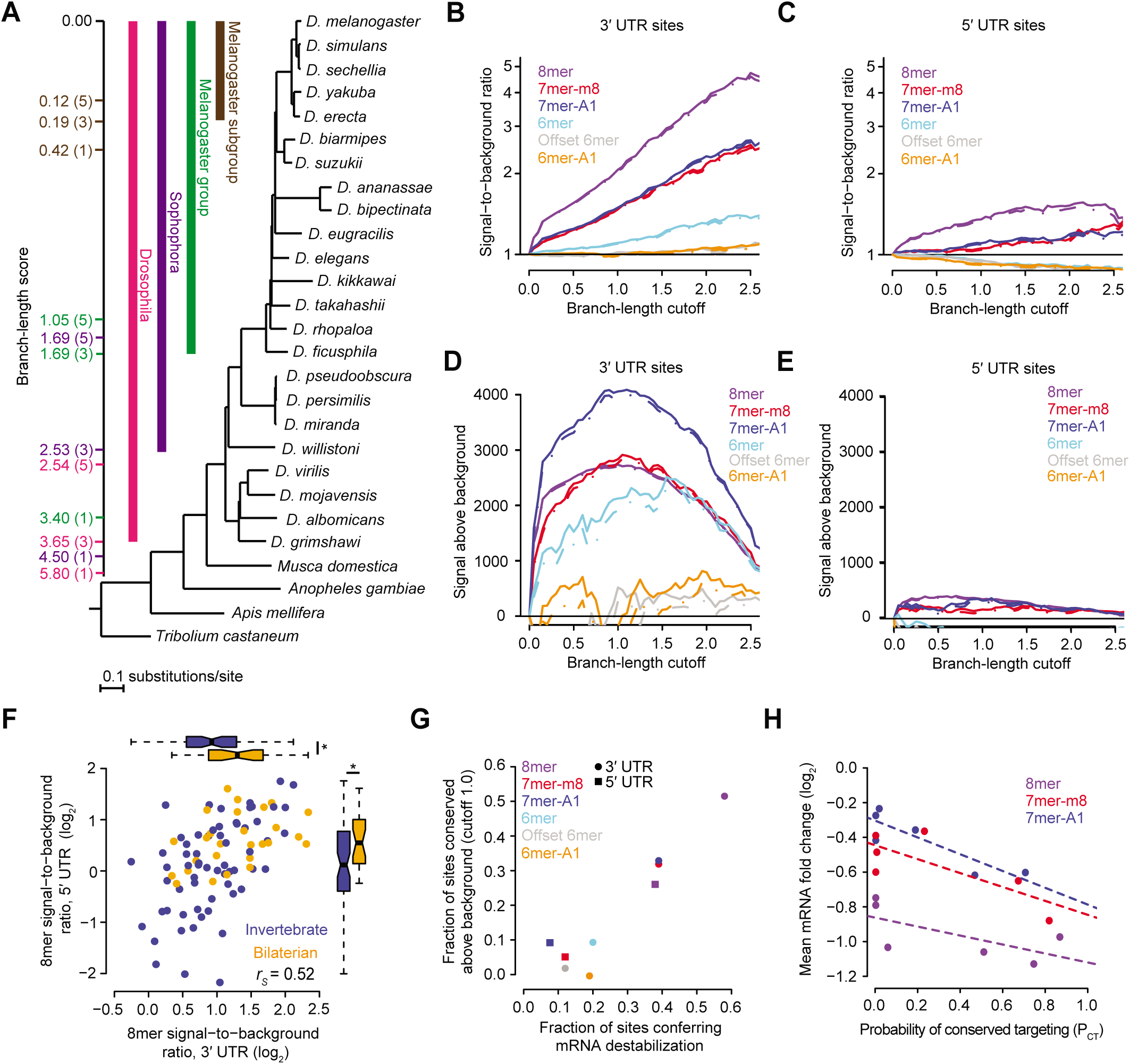
Evolutionary conservation of canonical sites in Drosophila 5′ UTRs and 3′ UTRs. **(A)** Phylogenetic tree of the 27 species used to examine miRNA site conservation. Outgroups of the genus *Drosophila* include *M. domestica* (the housefly), *A. gambiae* (the mosquito), *A. mellifera* (the European honey bee), and *T. castaneum* (the red flour beetle). *D. melanogaster* 3′ UTRs were assigned to one of five conservation bins based upon the median conservation of nucleotides across the entire 3′ UTR. The tree is drawn using the branch lengths and topology reported from genome-wide alignments in the UCSC genome browser. To the left of the tree, are colored coded branch-length scores corresponding to a site conserved among an entire subgroup of species indicated by a bar of the same color, showing scores for a site within a 3′ UTR in the lowest, middle, and highest conservation bins, labeled in parentheses as bins 1, 3, or 5, respectively. **(B–C)** Signal-to-background ratios for indicated site types at increasing branch-length cutoffs, computed for sites located in 3′ UTRs **(B)** or 5′ UTRs **(C)**. Broken lines indicate 5% lower confidence limits (z-test). These panels were modeled after the one originally shown for the analysis of mammalian 3′-UTR sites (Friedman et al. 2009). **(D–E)** Signal above background for indicated site types at increasing branch-length cutoffs, computed for sites located in 3′ UTRs **(D)** or 5′ UTRs **(E)**. Broken lines indicate 5% lower confidence limits (z-test). These panels were modeled after the one originally shown for the analysis of mammalian 3′-UTR sites (Friedman et al. 2009). **(F)** Signal-to-background ratios for the 8mer sites of 91 conserved miRNA seed families, calculated at near optimal sensitivity (a branch-length cutoff of 1.0), comparing the ratios observed for sites in 5′ UTRs to those for sites in 3′ UTRs (*r*s, Spearman correlation). Seed families conserved since the ancestor of bilaterian animals are distinguished from those that emerged more recently (orange and blue, respectively). Boxplots on the sides show the distributions of ratios for these two sets of families, with statistical significance for differences in these distributions evaluated using the one-sided Wilcoxon rank-sum test (\**P* < 0.01). See also Table S2. **(G)** Relationship between site conservation rate and repression efficacy. The fraction of sites conserved above background was calculated as ([Signal–Background]/Signal) at a branch-length cutoff of 1.0. The minimal fraction of sites conferring destabilization was determined from the cumulative distributions (Figures 1C and 1E), considering the maximal vertical displacement from the no-site distribution. Colors and shapes represent the canonical site types and UTR location, respectively. This panel was modeled after the one originally shown for the analysis of mammalian 3′-UTR sites (Friedman et al. 2009). **(H)** Relationship between site efficacy and site *P*CT. mRNAs were selected to have either one 7mer-A1, one 7mer-m8, or one 8mer 3′ UTR site to the transfected miRNA and no other canonical 3′-UTR site. mRNAs with sites of each type were grouped into six equal bins based on the site *P*CT. For each bin, mean mRNA fold change in the transfection data is plotted with respect to the mean *P*CT, with the dashed lines showing the least-squares fit to the data. The slopes for each are negative and significantly different from zero (*P* value < 10^−10^, linear regression using unbinned data).

For each site type of each of the 91 broadly conserved miRNA families, we computed the “signal” as the number of times that site occurred in *D. melanogaster* UTRs and had a branch-length score that equaled or surpassed a particular value (i.e., the “branch-length cutoff”). In parallel, we also computed the “background” as the number of conserved occurrences expected by chance, based upon the mean fraction of conserved motif instances for 50 length-matched *k*-mer controls, each of which was predicted to have background conservation resembling that of the miRNA site, as estimated from aggregated dinucleotide conservation rates (Friedman et al. 2009). This allowed us to compute a signal-to-background ratio at each branch-length cutoff, which represented the estimated enrichment of preferentially conserved miRNA sites in fly UTRs (Figure 2B and C). It also allowed us to compute the signal-above-background, which represented the estimated the number of miRNA sites that have been preferentially conserved in fly UTRs (Figure 2D and E).

As expected, the signal-to-background ratios increased as the evolutionary conservation criteria became more stringent, with 8mers in 3′ UTRs reaching a ratio of nearly five conserved sites for every one control site at the greater branch-length cutoffs (Figure 2B). For each site type, the ratios were consistently greater in the 3′ UTRs than they were in 5′ UTRs (Figure 2B and C). For example, in 5′ UTRs signal-to-background ratio for 8mers did not surpass 1.6 (Figure 2C). These results showed that sites are more likely to be conserved if they reside in 3′ UTRs, presumably because this is where they are also more effective (Figure 1). Nonetheless, when comparing the signal-to-background ratios for different miRNA families, ratios in 5′ UTRs correlated with those in 3′ UTRs (Figure 2F; Table S2). The greatest ratios tended to be for the fly miRNA families that have been conserved since the ancestor of bilaterian animals (Figure 2F), as might expected for these ancient families that have had more time to acquire broader roles in gene regulatory networks.

Although the sequence-conservation signal-to-background hierarchy of 8mer > 7mer > 6mer observed in both 5′ and 3′ UTRs matched the hierarchy observed for efficacy, some differences were observed. Most notably, the conservation signal for the 6mer site was robustly above background, whereas those for the offset 6mer and 6mer-A1 sites were both indistinguishable from background (Figure 2B), even though these three 6-nt sites had similar efficacies in our repression data (Figure 1C). Conversely, the 5′-UTR 7mer-A1 site exhibited a detectable signal for conservation (Figure 2B), even though it had no detectable efficacy in mediating repression (Figure 1C).

For sites in both 3′ and 5′ UTRs, the signal-above-background peaked near a branch-length cutoff of 1.0 (Figure 2D). At this and other branch-length cutoffs, the signal-above-background was far higher in the 3′ UTR than in the 5′ UTR (Figure 2D and E), which can be attributed to both a higher fraction of the sites preferentially conserved in 3′ UTRs, as indicated by the higher signal-to-background ratio in 3′ UTRs, and more sites residing in 3′ UTRs, mostly a consequence of 3′ UTRs generally being longer than 5′ UTRs. Including site types whose lower 5% confidence intervals exceeded zero, our results provided an estimate of ∼12,285 sites conserved above background in 3′ UTRs (2738 ± 31 8mer, 2837 ± 68 7mer-m8, 4062 ± 100 7mer-A1, 2128 ± 221 6mer sites, and 520 ± 244 offset 6mer sites, calculated at a branch-length cutoff of 1.0 and reported ± 90% confidence interval) (Figure 2D). When added to our estimate of ∼840 sites conserved above background in 5′ UTRs (350 ± 18 8mer, 165 ± 46 7mer-m8 sites, and 325 ± 44 7mer-A1 sites) (Figure 2E), the estimated number of preferentially conserved UTR sites in Drosophila UTRs totaled ∼13,125. Simulations that considered all of the conserved instances of site types and then accounted for those that were estimated to be conserved by chance in 5′ UTRs and 3′ UTRs, indicated that these 13,125 preferentially conserved sites reside within 5035 ± 83 (90% confidence interval) of the 13,550 unique mRNAs with annotated UTRs of Drosophila, implying that mRNAs from 37.2% ± 0.6% of the Drosophila genes are conserved targets of the broadly conserved miRNAs.

Additional comparison of the results from our analyses of site conservation and site efficacy revealed that, as observed for mammalian 3′-UTR sites (Friedman et al. 2009), there was a striking correlation between the fraction of sites conserved above background for each site type and the corresponding fraction of sites mediating mRNA destabilization (Figure 2G). Slightly deviating from this trend were 3′-UTR 6mer-A1 sites, which appeared to mediate some repression despite lacking a signal for conservation, and 5′-UTR 7mer-A1 sites, which had a modest signal for conservation despite undetectable efficacy of repression (Figure 2G).

To estimate the extent to which each instance of each of the three most effective sites has been preferentially conserved, we computed the probability of conserved targeting (*P*_CT_) score for each of the 8mer, 7mer-m8, and 7mer-A1 sites residing in *D. melanogaster* 3′-UTRs. *P*_CT_ scores, which range from 0 to 1, summarize the estimated probability that a given site has been evolutionarily conserved because of its pairing to the cognate miRNA, while controlling for other factors, such as its length, surrounding genomic context, and dinucleotide content (Friedman et al. 2009). These scores provide a valuable resource for biologists wanting to focus on conserved targeting interactions. They also can help predict targeting efficacy (Friedman et al. 2009; Agarwal et al. 2015). Indeed, sites with greater *P*_CT_ scores tended to confer more repression (Figure 2H), implying that as expected, conserved sites were more likely reside within contexts that favored their efficacy.

### Features useful for predicting site efficacy in flies

Before beginning to explore the features of site context associated with site efficacy, we improved the 3′-UTR annotations in S2 cells, the cell line in which we had acquired our functional data, reasoning that more accurate annotation of these UTRs would allow us to reduce the impact of false-positive sites while appropriately weighting sites by the frequency of their inclusion within 3′-UTR isoforms (Nam et al. 2014; Agarwal et al. 2015). Knowledge of abundant alternative 3′-UTR isoforms for the mRNAs of a gene would also provide a more informed assessment of 3′-UTR-related features, such as 3′-UTR length and distance from the closest 3′-UTR end. Accordingly, we identified and quantified the 3′-UTR isoforms of S2 cells using poly(A)-position profiling by sequencing (3P-seq) (Jan et al. 2011). Although the majority of the 3P-seq–supported poly(A) sites corresponded to either 3′-UTR isoforms that had been previously annotated by FlyBase or a large-scale study that annotated additional poly(A) sites (Smibert et al. 2012), nearly 47% of the 3P-seq–supported poly(A) sites did not correspond to existing annotations, and most of these novel sites could be linked to a nearby gene with the support of RNA-seq evidence (Figure 3A). In cases in which the longest 3′ UTR isoform for a gene annotated using 3P-seq differed from that annotated in FlyBase, it was more often longer, although for nearly 1000 genes the 3P-seq results implicated the dominant use of a shorter 3′-UTR isoform in S2 cells (Figure 3B). Using this information, we compiled a set of 3,826 mRNAs that passed our expression threshold in S2 cells and for which ≥90% of the 3P-seq tags corresponded to a single dominant 3′-UTR isoform in these cells, and used this set to investigate features of site context associated with site efficacy.

**Figure 3.**
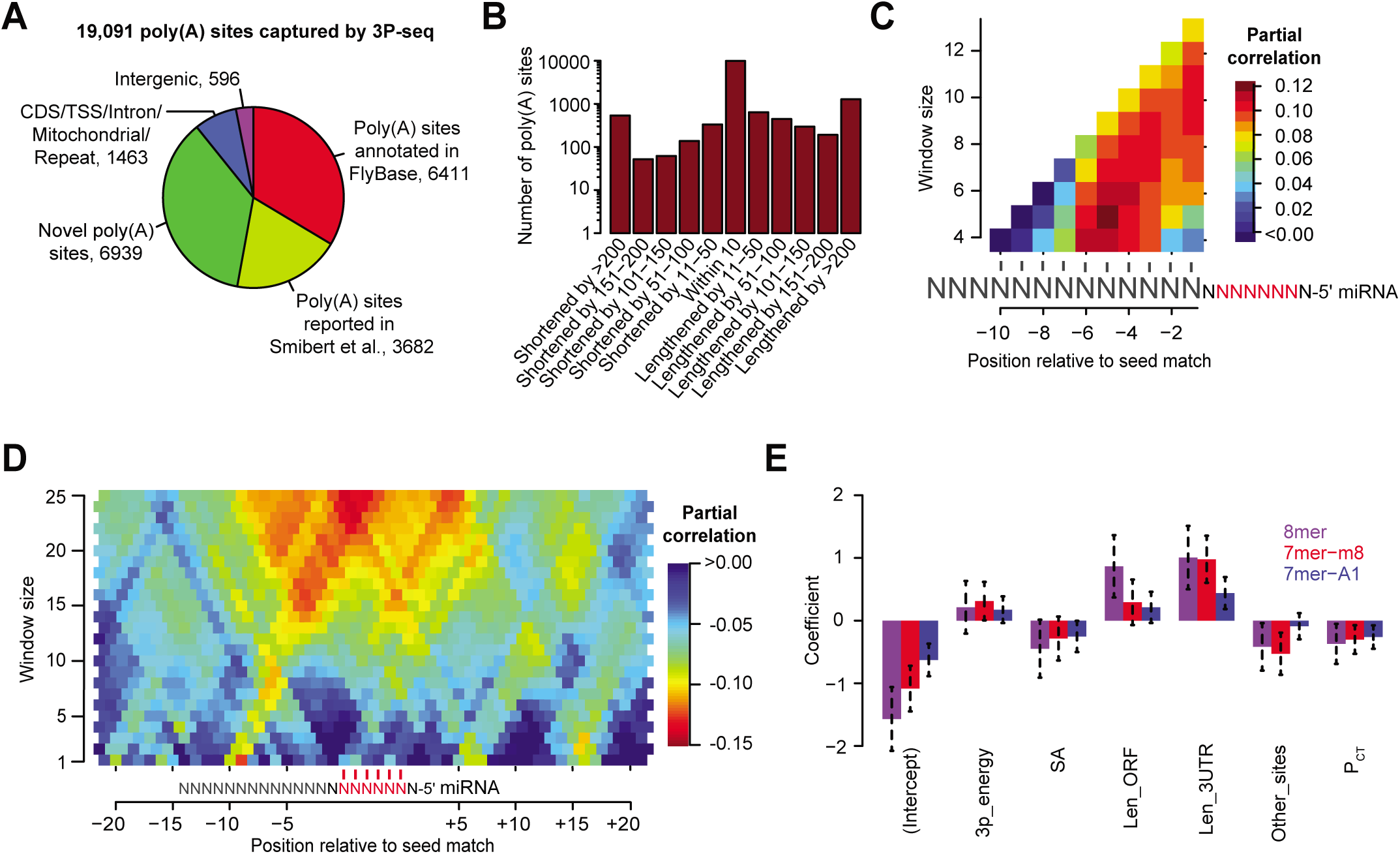
Refinement of 3′-UTR annotations in S2 cells and development of a regression model that predicts miRNA targeting efficacy in Drosophila. **(A)** Poly(A) sites detected in S2 cells by 3P-seq, classified with respect to their previous annotation status. **(B)** Extension and contraction of longest 3′-UTR isoforms relative to the FlyBase annotations. For each gene with a poly(A) site detected using 3P-seq, the difference between the longest 3′-UTR isoform annotated using 3P-seq was compared to longest 3′-UTR isoform annotated at FlyBase. These differences were then binned as indicated, and the number of sites assigned to each bin is plotted. **(C)** Optimization of scoring of predicted 3′-supplementary pairing in flies. Predicted thermodynamic energy scores were computed for the pairing between a 9-nt regions upstream of canonical 7–8-nt 3′-UTR sites and a variable-length region of the miRNA with the indicted size (window size) that began at the indicated position of the miRNA. The heatmap displays the partial correlations between these scores and the repression associated with the corresponding sites, determined while controlling for site type. **(D)** Optimization of the scoring of predicted structural accessibility in flies. Predicted RNA structural accessibility scores were computed as the average pairing probabilities for variable-length (window size) regions that centered at the indicated mRNA position, shown with respect to the seed match of each canonical 7–8-nt 3′-UTR site. The heatmap displays the partial correlations between these values and the repression associated with the corresponding sites, determined while controlling for site type. **(E)** The contributions of site type and each of the six features of the context model. For each site type, the coefficients for the multiple linear regression are plotted for each feature. Because features were each scored on a similar scale, the relative contribution of each feature in discriminating between more or less effective sites was roughly proportional to the absolute value of its coefficient. Also plotted are the intercepts, which roughly indicate the discriminatory power of site type. Bars indicate the 95% confidence intervals of each coefficient. See also Table S3, Table S4, and Figure S3.

With this set of mRNAs and repression values in hand, we examined two of the more complex features of site context, confirming their effects in Drosophila cells and developing scoring schemes that best correlated with their influence in these cells. The first of these two features was 3′-supplementary pairing, i.e., pairing to the target by miRNA nucleotides outside of the seed region. The strength of this pairing was evaluated as the predicted thermodynamic energy of pairing between the 3′ region of the miRNA and a corresponding mRNA region upstream of the seed match. This predicted energy of pairing was evaluated for mRNAs that possessed a single 7–8-nt 3′-UTR site for the transfected miRNA and then compared to the repression observed for the mRNAs upon miRNA transfection by computing a partial correlation between 3′-supplementary-pairing energies and mRNA changes, controlling for site type.

In mammalian cells, 3′-supplementary pairing is most influential when centered on nucleotides 13–17 (Grimson et al. 2007), but in flies the pairing possibilities most consequential for repression had not been identified. To systematically examine these possibilities, we varied three parameters: i) the start position of the miRNA region considered, examining all start possibilities from positions 9 to 19, ii) the length of the miRNA region considered, examining lengths from 4 to 13 nt, and iii) the length of the target region upstream of the seed match, examining lengths from 4 to 20 nt. A grid search over all parameter combinations revealed that the predicted energy of 3′-supplementary pairing energy was optimally predictive of repression efficacy when it was calculated for the pairing that can occur between miRNA nucleotides 13–17 and a 9-nt region upstream of the seed match (Figure 3C).

The second feature we investigated was the influence of 3′-UTR structure on target-site accessibility. This feature has been evaluated previously using two approaches, either evaluating nucleotide composition near the site, reasoning that sites residing in high local AU content would be more accessible (Grimson et al. 2007), or attempting to predict site accessibility using various RNA-folding algorithms (Robins et al. 2005; Kertesz et al. 2007; Hammell et al. 2008; Tafer et al. 2008; Hong et al. 2009; Agarwal et al. 2015). With respect to the second approach, a method originally developed to predict siRNA target-site accessibility (Tafer et al. 2008) appears to be one of the more effective methods for predicting miRNA target site accessibility in mammals (Agarwal et al. 2015). This method uses RNAplfold (Bernhart et al., 2006) to fold the 80-nt region centered on the seed match and then reports a structural accessibility (SA) score calculated as the mean unpaired probabilities for a smaller window in the vicinity of the seed match (Tafer et al. 2008; Agarwal et al. 2015). To determine optimal location and width of this window for scoring SA in flies, we again computed partial correlations, this time between mean pairing probabilities and mRNA changes, varying two parameters: i) the position of the center of the window within the target mRNA, examining each position within 20 nt of the seed match, and ii) the size of this widow, considering sizes of 1 to 25 nt. A grid search over all parameter combinations indicated that a 25-nt window centered on the nucleotide that pairs to miRNA position 7 was optimal for calculating SA in flies (Figure 3D). Although the optimal window size fell at the edge of the range, larger windows were not considered because they were more prone to extend beyond 3′-UTR boundaries, which reduced the sample size.

### A quantitative model for predicting site efficacy in flies

To identify and evaluate additional features associated with site efficacy in flies and generate a resource for placing fly miRNAs into gene regulatory networks, we developed a quantitative model of miRNA targeting efficacy for flies, which resembled our models developed for mammals (Grimson et al. 2007; Garcia et al. 2011; Agarwal et al. 2015). The smaller scope of our fly dataset imposed some limitations on the features we could examine in flies as well as the strategy used to train the model. In particular, the number of training examples was an order of magnitude lower in the fly dataset relative to the human dataset, which was due to i) fewer small-RNA transfection datasets in S2 cells compared to those available in HeLa cells, ii) a smaller number of genes expressed in S2 cells compared to those expressed in HeLa cells, and iii) shorter 3′ UTRs in flies, which further decreased the number of 3′ UTRs with a site for a miRNA of interest. Thus, we did not consider features related to the identity of the miRNA seed, such as target abundance, predicted seed pairing stability, and nucleotide identity at the miRNA or target position 8, which are each informative for predicting targeting efficacy in human cells (Garcia et al. 2011; Agarwal et al. 2015). Moreover, rather than considering features for each site type independently, we trained a single, unified regression model that considered the site type itself as a potential feature of targeting. In addition to site type, eight other features of the sites and their surrounding context and seven features of the target mRNAs were considered as potentially informative of targeting efficacy (Table 1).

**Table 1.** The 18 features considered in the models, highlighting the seven robustly selected through stepwise regression (bold). The feature description does not include the scaling performed (Table S2) to generate more comparable regression coefficients.

| Feature | Abbreviation | Description | Frequency chosen |
| --- | --- | --- | --- |
| **Site** |  |  |  |
| **Site type** | **site type** | **Type of site (8mer, 7mer-m8, or 7mer-A1) (Grimson et al., 2007)** | **100%** |
| Site position 1 | site1 | Identity of nucleotide at position 1 of the site | 0% |
| Site position 9 | site9 | Identity of nucleotide at position 9 of the site | 2% |
| Site position 10 | site10 | Identity of nucleotide at position 10 of the site | 0% |
| Local AU content | local_AU | AU content within 30 nucleotides of the site (Grimson et al., 2007) | 50% |
| 3'-supplementary pairing | 3P_score | Supplementary pairing at the miRNA 3' end (Grimson et al., 2007) | 4% |
| **Energy of 3'-supplementary pairing** | **3P_energy** | **Thermodynamic energy of supplementary pairing at the miRNA 3' end (dG duplex - dG seed duplex)** | **93%** |
| **Predicted structural accessibility** | **SA** | **log<sub>10</sub>(Probability that a 25-nt segment centered on the match to miRNA position 7 is unpaired)** | **91%** |
| **Probability of conserved targeting\*** | **P<sub>CT</sub>** | **Probability of site conservation, controlling for dinucleotide evolution and site context (Friedman et al., 2009)** | **100%** |
| **mRNA** |  |  |  |
| 5'-UTR length | len_5UTR | log<sub>10</sub>(Length of the 5' UTR) | 6% |
| **ORF length** | **len_ORF** | **log<sub>10</sub>(Length of the ORF) (Agarwal et al., 2015)** | **100%** |
| **3'-UTR length** | **len_3UTR** | **log<sub>10</sub>(Length of the 3' UTR) (Hausser et al., 2009)** | **100%** |
| 5'-UTR AU content | AU_5UTR | Fractional AU content in the 5' UTR | 21% |
| ORF AU content | AU_ORF | Fractional AU content in the ORF | 18% |
| 3'-UTR AU content | AU_3UTR | Fractional AU content in the 3' UTR | 51% |
| Distance from stop codon | dist_stop | log<sub>10</sub>(Distance of site from stop codon) | 4% |
| Minimum distance | min_dist | log<sub>10</sub>(Minimum distance of site from stop codon or poly(A) cleavage site) (Galdatzis et al., 2007; Grimson et al., 2007; Majoros and Ohler, 2007) | 57% |
| **Weak canonical sites in mRNA** | **other_sites** | **Number of 8mer sites in the 5' UTR and ORF and offset-6mer, 6mer-A1, and 6mer sites in the 3' UTR (Agarwal et al., 2015)** | **99%** |

\* Only relevant for deeply conserved miRNA families

Starting with these features, we trained models of targeting efficacy using a variety of machine-learning algorithms. To evaluate each algorithm, we partitioned our dataset into 1000 bootstrapped samples to estimate the held-out prediction performance. Each sample included 70% of the mRNAs with a single 7–8-nt 3′-UTR site from each miRNA transfection experiment (randomly selected without replacement), reserving the remaining 30% for testing. Among the different algorithms, a stepwise-regression strategy that maximized the Akaike Information Criterion (AIC) led to the best empirical performance (Figure S3). This stepwise-regression strategy was the same algorithm that we had recently used to build a model of mammalian miRNA targeting efficacy (Agarwal et al. 2015). Relative to a model that considered only site type (the “site only” model), the stepwise regression model that considered features of site context was 2–3 fold improved in predicting the mRNA fold-change measurements (median *r*^*2*^ of 0.08 and 0.19, respectively; p < 0.001, paired Wilcoxon signed-rank test; Figure S3). Note that because most of the variability in miRNA-transfection datasets is attributable to experimental noise and other factors that are not direct effects of miRNA targeting, the *r*^*2*^ of 0.19 in this analysis was not of concern and resembled the results obtained in mammalian analyses (Agarwal et al. 2015).

The features most informative for the stepwise regression model were presumably those with the greatest impact on site efficacy in flies. To identify these key features, we quantified the percentage of bootstrapped samples in which each feature was chosen (Table 1). Seven of the 18 features were selected in ≥ 90% of the bootstrap samples (Table 1), and multiple linear regression models trained with only these seven features performed at least was well as those that considered all 18 features (median *r*^*2*^ of 0.20; Figure S3). Aside from site type, which is currently considered in TargetScanFly, release 6.2 (Ruby et al. 2007), these robustly selected features included three features of the site, i.e, energy of 3′-supplementary pairing (3P_energy), structural accessibility (SA), and evolutionary conservation (*P*CT); and three features of the mRNA, i.e., ORF length (len_ORF), 3′-UTR length (len_3UTR), and the number of weak sites within the mRNA (other_sites) (Table 1). Notably, all of these features were previously selected when modeling site efficacy in mammals (Agarwal et al. 2015), with the nuance that in flies 3P_energy outperformed 3P_score, another method of evaluating 3′-supplementary pairing, which had been optimized on mammalian data (Grimson et al. 2007). However, two features strongly associated with site efficacy in mammals were not consistently selected in the fly analysis. These included AU composition in the vicinity of the target site (local_AU) and the minimum distance of a site from 3′-UTR boundaries (min_dist) (Grimson et al. 2007). Perhaps these features did not strongly discriminate effective targets from ineffective ones in flies because compared to mammalian 3′ UTRs, fly 3′ UTRs are constitutively more AU-rich and much shorter (median 3′-UTR length of 661 nt and 202 nt for human and fly, respectively, considering the longest UTR annotation per gene after removing genes with longest UTR annotations ≤ 2 nt).

Using the seven consistently selected features and the entire dataset of 3′ UTRs containing single 7mer-A1, 7mer-m8, or 8mer sites, we trained independent multiple linear regression models for each of these three canonical sites. These three models were then combined to generate a model for fly miRNA targeting, which we call the “context model” because it resembled our context models developed for mammalian miRNA targeting, in that it modeled site context in addition to site type. The sign of each coefficient revealed the relationship of each feature to repression (Figure 3E). For example, mRNAs with longer ORFs or longer 3′ UTRs, and sites with weaker 3′-supplementary pairing energy were more refractory to repression (as indicated by a positive coefficient), whereas target sites that were more structurally accessible or more conserved, and mRNAs with other weak sites were more prone to repression (as indicated by a negative coefficient). Normalizing the scores of each feature to a similar scale enabled assessment of the relative contribution of each feature to the context model (Figure 3E). As expected, site type was also a major predictor of repression in the model, as indicated by the large magnitude of the intercept term (Figure 3E). The signs and relative magnitudes of the features largely paralleled those found in the mammals (Agarwal et al. 2015), indicating that the influence of these features might reflect evolutionarily conserved aspects of miRNA targeting in bilaterian species. One difference was that *P*_CT_ scores contributed relatively more to the fly context model than they do to the analogous mammalian model (Agarwal et al. 2015), implying that the detection and scoring of the molecular features of target efficacy have more room for improvement in flies, presumably because less data were available in flies for feature identification and evaluation.

### Comparison to the performance of previous methods

We next compared the performance of the fly context model to that of previously reported methods, measuring how successfully each method predicted and ranked the mRNAs that respond to the gain or loss of a miRNA in Drosophila. For training, our context model had considered only mRNAs that had a single 7–8-nt site to the cognate miRNA within their 3′ UTR, but for testing it needed to be extended to mRNAs that had multiple sites to the same miRNA within their 3′ UTRs. Accordingly, for each predicted target, we generate a total context score, calculated as the sum the context scores of the sites to the cognate miRNA (Grimson et al. 2007), and used these total context scores to rank all of the predicted targets for each miRNA. The response of the top ranked targets was then compared to that of 12 previously reported methods, chosen because predictions for Drosophila targets were available online, as was information needed to rank these predictions. Having already generated the *P*_CT_ scores of the Drosophila sites, we also combined the scores of multiple 7–8-nt canonical sites when present within the same 3′ UTRs to generate Aggregate *P*_CT_ scores, which were then used to rank predictions based solely on the probability that they were preferentially conserved targets of the miRNA (Friedman et al. 2009).

We took precautions to perform a fair comparison of the algorithms. First, for each algorithm, we considered only predicted targets that corresponded to mRNAs expressed above the quantification threshold in the relevant test-set sample lacking the miRNA. Second, we avoided testing the context model on the same transfection data upon which it was trained. More specifically, we implemented a cross-validation strategy when testing the results of the context model using the transfection datasets, sequentially holding out each dataset and re-training the coefficients for the features in our context model using the five remaining transfection datasets before generating predictions for the held-out dataset. Further reducing the concern of overfitting was the observation that most top-ranked targets contained two or more canonical 3′ UTR sites and thus were not used during the development and training of our model. Third, for all testing of the context model, we used coefficients retrained on publicly available FlyBase 3′-UTR annotations, reasoning that training on improved 3′-UTR annotations derived from our 3P-seq data would have imparted an advantage to our model.

Testing was initially performed using our six datasets that each examined mRNA changes after transfecting a miRNA into S2 cells. For each algorithm and each transfected miRNA, we computed the mean mRNA fold change of the top-ranked targets of the transfected miRNA and then plotted the mean value for the six different miRNAs at various ranking thresholds, thereby summarizing repression efficacy of the top-ranked targets at each threshold (Figure 4A). Some methods, such as PicTar, which generated relatively few predictions, could be evaluated at only a few thresholds, whereas others, such as RNA22 and TargetSpy, could be evaluated at many more (Figure 4A). With the exception of RNA22, all algorithms predicted repressed targets better than expected by chance. However, some, including PicTar, MinoTar, RNAhybrid, TargetSpy, and mirSVR, performed similarly or worse than a naïve strategy of selecting all mRNAs that have at least one 7–8-nt canonical site in their 3′ UTR. Of the previously reported algorithms, TargetScanFly, EMBL, and PITA.Top performed the best. Nevertheless, our context model performed better than all previous methods, providing predictions that were the most responsive to transfection of the miRNA at each threshold tested (Figure 4A).

**Figure 4.**
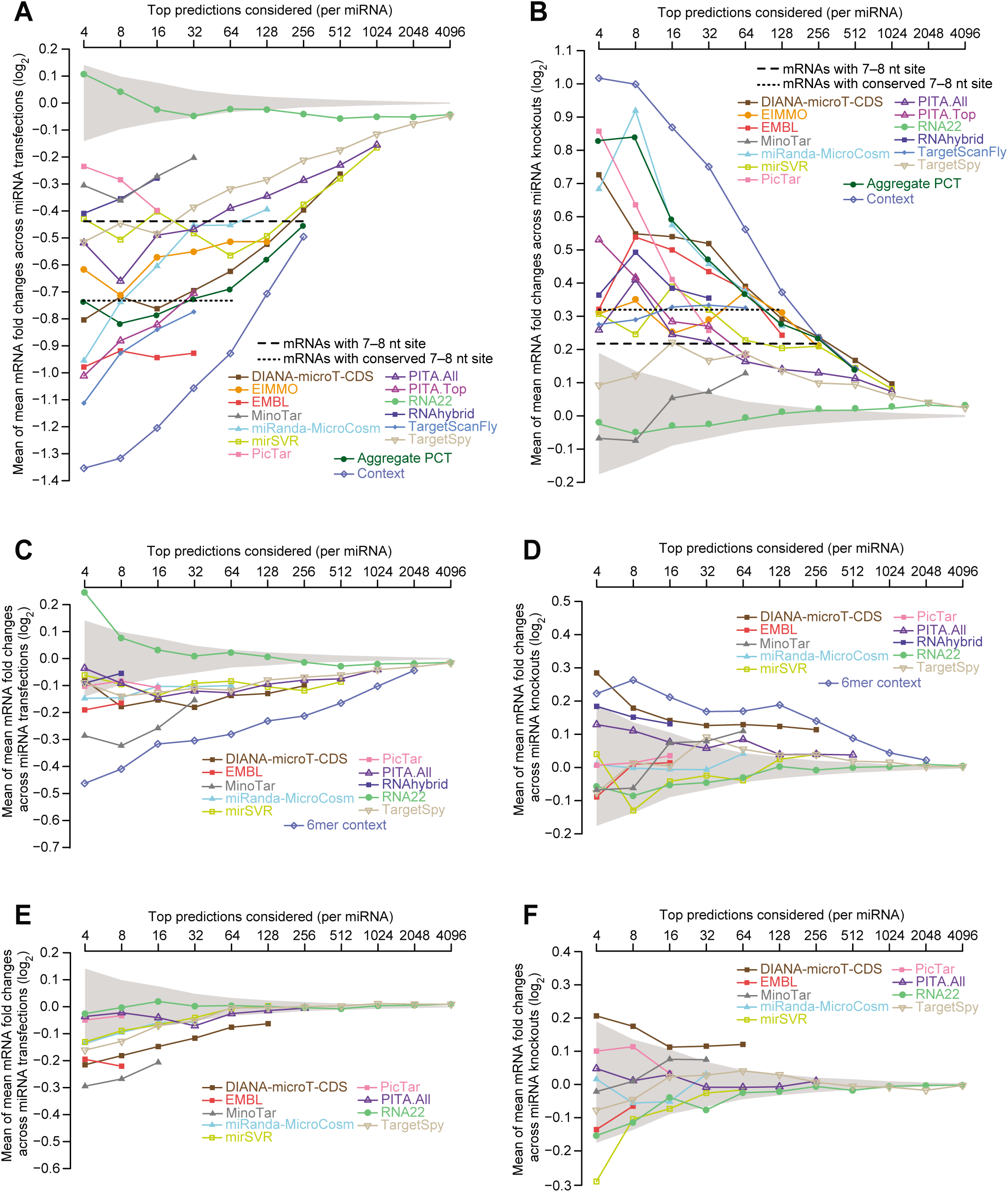
Performances of different target-prediction algorithms in flies. **(A)** The differential ability of algorithms to predict the mRNAs most responsive to miRNAs transfected into Drosophila cells. Shown for each algorithm in the key are mean mRNA fold changes observed for top-ranked predicted targets, evaluated over a sliding sensitivity threshold using the six miRNA transfection datasets. For each algorithm, predictions for each of the six miRNAs were ranked according to their scores, and the mean of the mean fold-change values was plotted at each sensitivity threshold. For example, at a threshold of 16, the 16 top predictions for each miRNA were identified (not considering predictions for mRNAs expressed too low to be accurately quantified). mRNA fold-change values for these predictions were collected from the cognate transfections, and the mean fold-change values were computed for each transfection for which the threshold did not exceed the number of reported predictions. The mean of the available mean values was then plotted. Also plotted are the mean of mean mRNA fold changes for all mRNAs with at least one cognate canonical 7–8-nt site in their 3′ UTR (dashed line), the mean of mean fold change for all mRNAs with at least one conserved cognate canonical 7–8-nt site in their 3′ UTR (dotted line), and the 95% confidence interval for the mean of mean fold changes of randomly selected mRNAs, determined using 1000 resamplings (without replacement) at each cutoff (shading). Sites were considered conserved if their branch-length scores exceeded a cutoff with a signal:background ratio of 2:1 for the corresponding site type (cutoffs of 1.0, 1.6, and 1.6 for 8mer, 7mer-m8, and 7mer-A1 sites, respectively; Figure 2B). See also Figure S4. **(B)** The differential ability of algorithms to predict the mRNAs most responsive to knocking out out miRNAs in flies. Shown for each algorithm in the key are mean mRNA fold changes observed for top-ranked predicted targets, evaluated over a sliding sensitivity threshold using the three knockout datasets. Otherwise, this panel is as in (A). **(C and D)** The differential ability of algorithms to predict targets that respond to the miRNA despite lacking a canonical 7–8-nt 3′-UTR site. These panels are as in (A) and (B), except they plot results for only the predicted targets that lack a canonical 7–8-nt site in their 3′ UTR. Results for our context model and other algorithms that only predict targets with canonical 7–8-nt 3′-UTR sites are not shown. Instead, results are shown for a 6mer context model, which considers only the additive effects of 6mer, offset 6mer, and 6mer-A1 sites and their corresponding context features. **(E and F)** The difficulty of predicting mRNAs that respond to miRNA transfection or knock out despite lacking canonical 6–8-nt 3′-UTR sites. These panels are as in (C and D), except they plot results for mRNAs with 3′ UTRs that lack a canonical 6–8-nt site.

Although our cross-validation strategy avoided testing our model on the same measurements as used for its training, some concerns regarding testing on the transfection data remained because these data was used to optimize scoring of some features of our model. Moreover, transfection introduces high concentrations of miRNAs to in cells in which they normally are not acting, raising the concern that a model developed and tested solely on transfection datasets might not accurately predict the response of miRNAs in their endogenous physiological contexts. Therefore we searched for a test set that had not been used to develop any of the algorithms and that monitored the transcriptome response to endogenous miRNAs expressed at physiological levels. Surveying the Drosophila literature, we identified three miRNA-knockout datasets with compelling signals for miRNA-mediated repression. Pooling these datasets, which monitored mRNA changes after deleting either miR-14 (Varghese et al. 2010), miR-34 (Liu et al. 2012), or miR-277 (Esslinger et al. 2013), and carrying out the same type of analysis as we had done for the transfection datasets (but monitoring de-repression following loss of a miRNA instead of repression following introduction of a miRNA) revealed performances that generally resembled those observed with the transfection datasets (Figure 4B). The relative performances of the previous methods shifted somewhat, with improvement observed for Aggregate *P*CT, miRanda-MicroCosm, and PicTar and worsening observed for MinoTar, TargetScanFly, and TargetSpy. Importantly, however, when testing on these consequences of endogenous miRNA targeting in flies, the context model again performed better than all previous models. Results for miR-277 resembled those for the other two miRNA (data not shown), even though miR-277 is unusual in that it primarily resides within Ago2 rather than Ago1 (Forstemann et al. 2007).

The mean-of-means metric that we used to evaluate repression of top-ranked targets had several potential limitations. For example, it can exaggerate the influence of individual outliers, or more heavily weight datasets with a greater variance in their fold-change distributions. Nonetheless, examination of plots showing the mean of median mRNA changes did not substantially change our assessment of the relative performance of each algorithm, which indicated that we did not arrive at erroneous conclusions because of outliers (Figure S4). Also potentially influencing our comparisons was the fact that for some previous algorithms predictions were missing for some miRNAs of our test sets. For example, EMBL predictions were not available for miR-263a and miR-994, and because targets for these two miRNAs happened to undergo less repression in our transfections, the testing of EMBL on only the remainder of the transfection datasets presumably inflated its relative performance.

Target-prediction algorithms have been developed with divergent priorities regarding prediction accuracy. Out of concern for prediction specificity, some, including our context model, consider only predictions with the most effective types of sites, i.e., 7–8-nt seed-matched sites within 3′ UTRs. In contrast, other algorithms, out of concern for prediction sensitivity, do not limit their predictions to those with these most effective site types, and some of these include predictions with a vast array of non-canonical sites that show no evidence of efficacy when tested using data from mammals and fish (Agarwal et al. 2015). To begin to explore the tradeoffs of these divergent priorities when predicting miRNA targets in flies, we removed predictions containing 7–8-nt canonical sites to the cognate miRNA in their 3′ UTRs, and tested the behavior of the remaining predictions that lack these more effective canonical sites. When testing on the transfection data, most algorithms that do not strictly focus on 3′ UTRs with 7–8-nt canonical sites generated predictions that were repressed more than expected by chance (Figure 4C).

Encouraged by these results, we used our context features to build a model that considered predictions that lacked canonical 7–8-nt 3′-UTR sites but had at least one offset 6mer, 6mer, or 6mer-A1 site in their 3′ UTR. When using either test set and testing only predictions that lacked canonical 7–8-nt 3′-UTR sites to the cognate miRNA, this model, which we call the “6mer context” model, performed better than all existing algorithms (Figure 4C and D). The other algorithm that yielded predictions consistently repressed better than background was DIANA-microT-CDS, which includes predictions with only canonical ORF sites. Thus, taken together, our analysis indicates that two distinct strategies that focus on only marginally effective sites can be predictive in flies, as judged by both transfection and knockout results; one approach focuses on canonical 6-nt sites in 3′ UTRs, and the other focuses on canonical ORF sites. However, at best, the average repression of the 4–8 top predictions from these approaches was much less than that of the top targets of the standard context model and instead resembled that of the hundreds of mRNAs that contained 7–8-nt canonical 3′-UTR sites (Figure 4A–D).

The observation that models could be built that successfully predicted targets with only marginal canonical sites was consistent with the demonstrated efficacy of these marginal sites in Drosophila cells (Figure 1). A larger challenge has been to predict effective non-canonical sites, which lack at least a 6-nt perfect match to the seed region. Although two types of non-canonical sites, known as the 3′-supplementary sites and centered sites, can mediate repression, these sites are rare—indeed so rare that is difficult to observe a signal for their action in mammalian cells without aggregating many datasets (Bartel 2009; Shin et al. 2010). Nonetheless, some algorithms yield many predictions that have only non-canonical sites. Analyses of mammalian datasets indicate that these predictions are no more repressed than expected by chance (Agarwal et al. 2015), raising the question as to whether any of the algorithms might successfully predict non-canonical sites in Drosophila. To answer this question, we used our two test sets to measure the response of predictions that lacked any canonical 6–8-nt site to the cognate miRNA in their 3′-UTR (Figure 5E–F). The only predictions with a convincing signal above background in either test set were those of EMBL, DIANA-microT-CDS, and MinoTar. Manually examining the top-ranked predictions from EMBL revealed that the signal observed for its predictions was attributable to canonical sites located in ORFs and 3′ UTRs of alternative last exons, whereas the signal for the predictions of DIANA-microT-CDS and MinoTar was attributable canonical ORF sites. We conclude that in flies, as in mammals (Agarwal et al. 2015), non-canonical sites only rarely mediate repression, although we cannot exclude the formal possibility that effective non-canonical sites are abundant yet for some reason not predicted above background by any of the existing algorithms.

Having found that the context model performed better than the models that have been providing target predictions to the Drosophila research community (Figure 4A–B), we have set out to overhaul TargetScanFly (available at targetscan.org) to display these predictions. This new version of TargetScanFly (v7.0) will provide any biologist with an interest in either a miRNA or a potential miRNA target convenient access to the predictions, with an option of downloading code or bulk output suitable for more global analyses. Because of the diminishing returns of predicting targets with only marginal sites (Figure 4C–F), TargetScanFly will continue to focus on predictions with 7–8-nt canonical 3′-UTR sites, with ranks driven by the version of the context model that was trained on the entire transfection dataset. We have also released the accompanying TargetScanTools (https://github.com/vagarwal87/TargetScanTools) to help others reproduce our analyses, apply our computational pipeline to future datasets, and produce figures analogous to those of this manuscript. With these new resources, we hope to enhance the productivity of miRNA research in flies and thereby accelerate the understanding of this intriguing class of regulatory RNAs.

## METHODS

### Cell culture

*Drosophila* Schneider 2 (S2) cells were grown in Express Five serum-free media (GIBCO) supplemented with glutamine to 16 mM. Upon reaching confluency (every ∼ 3-5 days), cells were passaged following mechanical resuspension with a scraper (Corning). Prior to resuspension, the media and any unattached cells were removed and replaced with an equal volume of fresh media in order to select for attached cells.

### MicroRNA transfection, FACS, and mRNA isolation

Prior to transfection, cells were seeded into 6-well plates (Corning) at 2.5 x 10^6^ cells and 2 ml media per well. After 24 hours, each well was co-transfected with 2.5 μg plasmid (25% p2032-GFP, 75% pUC19) plus 25 nM miRNA duplex (or for mock transfections, with plasmid only) using 5 μl DharmaFECT Duo (Dharmacon). Equal volumes of nucleic acid and DharmaFECT Duo diluted in 1X PBS were combined and incubated at room temperature for 20 minutes to form transfection complexes that were then added dropwise to the cells (500 μl/well). Twenty-four hours after transfection, cells were harvested, resuspended in 1X PBS, passed through a 70 μm filter, and stained with 5 μg/ml propidium iodide (PI). For each transfection, 3–5 x 10^6^ GFP-positive and PI-negative cells were isolated by FACS and lysed in 1 ml TRI Reagent (Ambion). Following extraction from the lysate, total RNA was cleaned up using the RNeasy Mini kit (Qiagen) and subjected to poly(A) selection using oligo(dT) Dynabeads (Invitrogen) to isolate mRNA.

### Preparation of sequencing libraries

Strand-specific mRNA-Seq libraries for Illumina sequencing were prepared as described (Guo et al. 2010), with differences noted below. Briefly, poly(A)-selected RNA was hydrolyzed in alkaline buffer, resulting in fragments bearing 5′-hydroxyl and 3′-phosphate groups. Fragments between 36–55 nt were size selected and end-specific adapters were sequentially ligated onto each terminus; prior to each ligation step, the appropriate 3′ or 5′ end chemistry was generated through dephosphorylation or phosphorylation, respectively. Adapter-flanked fragments were reverse transcribed and the resulting cDNA PCR-amplified using primers complementary to each adapter. The PCR products were purified on a denaturing formamide gel and submitted for deep sequencing. 3P-seq libraries were prepared from RNA isolated from S2 cells as described (Jan et al. 2011).

### RNA-seq analysis

RNA-seq reads were analyzed using the quantification pipeline previously described (Denzler et al. 2014; Wong et al. 2015). A genome index was built for the latest build of the *D. melanogaster* genome (dm6) using STAR v. 2.4 (options --runMode genomeGenerate-- genomeFastaFiles dm6.fa-- sjdbGTFfile dmel-all-r6.07.gff-sjdbOverhang 40-- sjdbGTFtagExonParentTranscript Parent) (Dobin et al. 2013), with “dmel-all-r6.07.gff” referring to fly transcript models annotated in FlyBase release 6.07 (dos Santos et al. 2015), processed to have a single “Parent ID/exon” combination per line. Raw reads were aligned to the index with STAR (options --outFilterType BySJout --outFilterMultimapScoreRange 0 --readMatesLengthsIn Equal --outFilterIntronMotifs RemoveNoncanonicalUnannotated-clip3pAdapterSeq TCGTATGCCGTCTTCTGCTTG --outSAMstrandField intronMotif --outStd SAM). Considering all replicates of a particular sample, mRNA fold changes were computed between the miRNA transfection library of interest and the three mock-transfection biological replicates, using cuffdiff v. 2.2.1 (options --library-type fr-secondstrand-b dm6.fa-u --max-bundle-frags 100000000) (Trapnell et al. 2013), using protein-coding genes gene models from FlyBase release 6.07 (dos Santos et al. 2015).

### Selection of mRNAs for computational analysis

To avoid noisy mRNA fold-change measurements of poorly expressed genes, we used only genes whose expression values (measured in Fragments Per Kilobase Million, FPKM) exceeded 5.0 in the mock condition for all subsequent analyses. This threshold was chosen based upon visual inspection of plots evaluating the relationship between mean expression level and fold change (commonly known as “MA plots” in the context of microarrays), attempting to balance the tradeoff between sample size and noise reduction. To select gene annotations for site efficacy, data normalization, and evolutionary analyses (i.e., for Figure 1, Figure S1, and Figure 2, respectively), we selected one representative transcript isoform per gene, choosing the transcript isoform with the longest ORF, and if tied, the one with the longest 3′ UTR, and if still tied, the one with the longest 5′ UTR. This representative transcript was supplemented with the longest 3′ UTR among the subset of transcripts that shared the same stop codon.

To select gene annotations for feature optimization and regression modeling (i.e., for Figure 3 and Figure S3), we analyzed 3P-seq data to quantify the relative abundance of 3′-UTR isoforms related to each representative transcript. We then selected the subset of mRNAs for which ≥90% of the 3P-seq tags corresponded to a single dominant 3′-UTR isoform and used this dominant 3′-UTR isoform as the annotation for the corresponding gene. These steps followed the training framework previously described (Agarwal et al. 2015).

To select gene annotations for evaluation of model performance (i.e., for Figure 4 and Figure S4), we identified the longest and shortest 3′-UTR isoforms, as annotated by FlyBase, corresponding to each representative transcript. Context scores and aggregate *P*_CT_ scores were then generated for the longest and shortest 3′-UTR isoform groups separately, and then, for each gene and miRNA combination, the scores were averaged between the longest and shortest isoforms. To filter out targets with a predicted target site (i.e., for Figure 4B/D and Figure S4), we removed those that contained the relevant site types in the 3′ UTR of their representative transcript.

### Dataset normalization

mRNA changes correlated among the six transfection experiments, indicating the presence of batch effects and other biases (Figure S1A). To remove biases in the mRNA fold-change measurements, we implemented our previously described normalization strategy (Agarwal et al. 2015), which uses partial least-squares regression (PLSR) to remove sources of variation that are common to multiple independent miRNA transfections. This led to a modest improvement in our ability to detect signatures of miRNA-mediated target repression (Figure S1B–D). However, 5′-UTR length, ORF length, 3′-UTR length, 5′-UTR AU content, ORF AU content, 3′-UTR AU content, and mock-transfection gene expression level still correlated with fold changes for mRNAs with no predicted miRNA target site. The magnitude of these correlations varied significantly when comparing results of different miRNA transfection experiments. Thus, for each of the six miRNA transfection experiments, we fit a multiple linear regression model between the mRNA fold changes (i.e., which had already been normalized by the PLSR model) and the seven aforementioned features, using log-transformed values for the expression level feature. Although only mRNAs with no predicted canonical miRNA target site were used for this fit, the resulting linear model was used to predict mRNA fold changes for all mRNAs (including those with a predicted site), and for each gene, the residual value (the difference between the mRNA fold change and predicted mRNA fold change) was designated as its final normalized mRNA fold change (Table 1). Applying this second normalization to data from each transfection experiment led to enhanced detection of target repression, as indicated by a shift towards more significant p-values, especially for mRNAs with 3′ UTRs that contained weaker site types (Figure S1D).

Each miRNA transfection exhibited a variable level of global target repression (Figure S2). Reasons for this variability presumably included variability in transfection efficiency and differences in either the target abundance (TA) or the predicted seed pairing stability (SPS) of the miRNAs tested (Garcia et al. 2011; Agarwal et al. 2015). Because we did not have the power in sample size to accurately model the effects of either SPS or TA, as was possible in mammals (Garcia et al. 2011; Agarwal et al. 2015), we normalized the transfections to the same scale prior to training and testing the model. To do so, for each transfection dataset *D*, we computed the upper and lower quartiles of the mRNA log fold changes (*UQD* and *LQD*, respectively) as well as the corresponding quartiles for the fold changes among all datasets pooled together (*UQP* and *LQP*). We then updated each fold change *x* as follows: 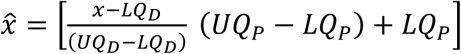. By centering on quartiles, this procedure normalized the fold-change distributions in a way that was less susceptible to the influence of outliers.

### Refining 3′-UTR isoform annotations

3P-seq data were processed as previously described (Ulitsky et al. 2012) but with adjustment of some of the parameters to better fit the characteristics of the fly 3′ UTRs. Transcript models were identified using Cufflinks and the ModENCODE S2 RNA-seq data (SRA accession SRR070279) with default parameters and minimum intron length set to 10. 3P-seq reads were processed and aligned to the dm3 genome assembly as described (Jan et al. 2011) and the resulting tag positions were lifted over to the dm6 assembly using the UCSC liftOver tool. In the first step of 3′UTR annotation, clusters of 3P-seq tags were generated as described (Ulitsky et al. 2012). Briefly, positions were sorted in descending order based on read count and the list was traversed such that for the position with the highest read count (or the first encountered read, in the case of a tie), all the tags within 30 nt were grouped and removed from the list as a cluster. Each cluster represented by a position with at least three total reads and at least two unique reads was considered a poly(A) site and was assigned the representative position supported by the most reads. RNA-seq data were then used to test if the poly(A) site connected with transcript models, as described previously (Ulitsky et al. 2012). Connectivity to gene models was established based on the Cufflinks gene models, allowing for gaps up to 200 nt. 3′ UTRs ending within 30 nt of each other were grouped together and assigned with their combined read count. The longest 3′ UTR of a gene was one with the maximal exonic length and which accounted for at least 1% of the 3P-seq reads. Other parameters were as described before (Ulitsky et al. 2012). A poly(A) site was considered to be “known” if it mapped within 20nt of a FlyBase poly(A) site. 3p-seq tags mapped to the dm6 genome, processed into clusters, and annotated can be found as BED files associated with Figure 2 at https://github.com/vagarwal87/TargetScanTools.

### MicroRNA sets

All mature fly miRNAs were downloaded from miRBase release 21 (Kozomara and Griffiths-Jones 2014). Those that matched a conserved miRNA at nucleotides 2–8 were considered part of that miRNA family. When partitioning miRNA families according to their conservation level, we compared the previously defined set of conserved families available in TargetScanFly (Ruby et al. 2007) with a more recent annotation of conserved “Pan-Drosophilid” miRNA families (Mohammed et al. 2013). For each difference between the two sets, we compared whether nucleotides 2–8 of each miRNA were conserved among most Drosopholids beyond the Sophophoran clade, as determined from the 27-way multiz alignments of each mature miRNA from the UCSC Genome Browser (Blanchette et al. 2004; Karolchik et al. 2014). This filter led to the removal of several miRNAs from being considered broadly conserved (e.g., mir-307b, mir-973, mir-975, mir-1014, mir-4977, and mir-4987) and the choice of a set of 91 conserved miRNA families (Table S2). From these 91, the set of 28 families conserved since the ancestor of bilaterian animals was identified, starting with a previous annotation of bilaterian miRNA families (Fromm et al. 2015), but separating related bilaterian families with different seed sequences and requiring that for each family the ancestral seed sequence has been conserved to Drosophila without a substitution or a shift in register (Table S2).

All miRNAs annotated in miRBase but not meeting our criteria for conservation were also grouped into families based on the identity of nucleotides 2–8 and were classified as poorly conserved miRNAs (which included many small RNAs misclassified as miRNAs). These miRNA seed families will be available for download at TargetScanFly (targetscan.org).

### Evolutionary analyses and calculation of *P*_CT_ scores

Fly *P*_CT_ scores were computed using the following datasets: i) 5′ UTRs or 3′ UTRs derived from 13,454 fly protein-coding genes annotated in FlyBase 6.07 (dos Santos et al. 2015), and ii) regions of multiple sequence alignments corresponding to these 5′-or 3′-UTRs, derived from the 27-way multiz alignments of the insect clade in the UCSC genome browser, which used the *D. melanogaster* genome release dm6 as its reference species (Blanchette et al. 2004; Karolchik et al. 2014). We partitioned 5′ UTRs and 3′ UTRs into five conservation bins based upon the median branch-length score (BLS) of the reference-species nucleotides, following the strategy previously described (Friedman et al. 2009; Jan et al. 2011). BLSs were computed using the BranchLengthScoring.py script from MotifMap (Daily et al. 2011). We used an updated computational pipeline for evolutionary analysis described previously (Agarwal et al. 2015) to estimate branch lengths of the phylogenetic trees for each bin, to compute the rates of *k*-mer conservation for canonical sites and control *k-*mers, and to calculate *P*_CT_ parameters and scores. All phylogenetic trees and *P*_CT_ parameters will be available for download at the TargetScanFly website (targetscan.org).

### Estimating the number of genes with preferentially conserved sites

A simulation was performed to estimate the number of genes containing a conserved site after accounting for the background of conserved sites. Towards this goal, we first identified for each conserved miRNA all unique target sites with BLS ≥ 1.0, yielding a total of 8,743 5′-UTR sites (considering 8mer, 7mer-m8, and 7mer-A1 sites) and 86,872 3′-UTR sites (considering 8mer, 7mer-m8, 7mer-A1, 6mer sites, and offset 6mer sites) that surpassed this cutoff. Among these, we estimated that 840 ± 40 5′-UTR sites and 12,285 ± 214 3′-UTR sites (mean ± standard deviation) were conserved above background. To estimate the distribution of genes with conserved sites, we performed 1000 samplings with the following procedure: i) An integer was randomly selected from each of the two normal distributions of total sites above background. ii) Using each of these two integers, a corresponding number of conserved sites was randomly sampled (without replacement) from the respective 5′ UTRs or 3′ UTRs. iii) The number of unique genes containing the selected sites was recorded. After 1000 samplings, the distribution of values obtained for our estimate of genes with conserved sites had a mean of 5035 and a 90% confidence interval of ± 83.

### Regression models

3P_energy was scored as described in the text. Other features were scored was as described (Agarwal et al. 2015), except SA was scored used the parameters optimized for Drosophila. For each feature of the final context model, scores were scaled (Table S3) before being multiplied by their corresponding coefficients (Table S4).

To evaluate performance, we generated 1000 bootstrap samples in which we used, for each site type and transfection experiment, 70% of data to train the models and the remaining data as a test set. To choose a model, we evaluated the performance of a variety of machine-learning strategies, including i) “all subsets regression”, maximizing the Bayesian information criterion (BIC) as implemented in the *regsubsets* function of the “leaps” R package (parameters “nvmax=15, nbest=1, method = ‘forward’, really.big=T”), ii) stepwise regression, maximizing the BIC or Akaike information criterion (AIC) as implemented in the *stepAIC* function from the “MASS” R package (Venables and Ripley 2002), iii) Lasso regression using the *cv.glmnet* function (parameters “nfolds=10, alpha=1”) in the “glmnet” R package, iv) Multivariate Adaptive Regression Splines (MARS) as implemented in the “earth” R package (parameters “degree = 1, trace = 0, nk = 500”), v) random forest regression using the “randomForest” R package, vi) Principal Components Regression (PCR) or Partial Least Squares Regression (PLSR) using the *pcr* and *plsr* functions as implemented in the “pls” R package (parameter “ncomp = 5” during prediction). As for our model of mammalian targeting (Agarwal et al. 2015), we ultimately utilized stepwise regression, with AIC to select features.

For the model driving TargetScanFly, we fit a multiple linear regression model for each site type using the selected group of features, training with all of the genes that were expressed above the threshold in our transfection datasets and had single 3′-UTR sites and 90% UTR homogeneity. As for mammalian predictions (Agarwal et al. 2015), scores for 8mer, 7mer-m8, and 7mer-A1 sites were bounded to be no greater than - 0.03, - 0.02, and - 0.01, respectively, thereby creating a piecewise linear function for each site type. For each 3′ UTR with at least one 7–8-nt site to the miRNA, the context scores of the sites were summed together to acquire a total context score, which was used to rank the predicted target gene (Figure 4 and Figure S4).

### Performance comparisons

To compare predictions from different miRNA target prediction tools, we collected the following downloadable predictions: DIANA-microT-CDS (September 2013) (Reczko et al. 2012), EIMMo v5 (January 2011) (Gaidatzis et al. 2007), EMBL (2005 predictions) (Brennecke et al. 2005; Stark et al. 2005), miRanda-MicroCosm v5 (Griffiths-Jones et al. 2008), mirSVR (August 2010) (Betel et al. 2010), PicTar (from the doRina web resource; sets conserved among D. melanogaster, D. yakuba, D. annanasae, D. pseudoobscura, D. mojavensis, and D. virilis) (Grun et al. 2005; Anders et al. 2012), PITA Catalog v6 (3/15 flank for either “All” or “Top” predictions, August 2008) (Kertesz et al. 2007), RNA22 (May 2011) (Miranda et al. 2006), RNAhybrid (Rehmsmeier et al. 2004), TargetSpy (all predictions) (Sturm et al. 2010), MinoTar (downloaded from TargetScanFly ORF v6.2, June 2012) (Schnall-Levin et al. 2010), and TargetScanFly v6.2 (June 2012) (Ruby et al. 2007). For algorithms providing site-level predictions (i.e., ElMMo, mirSVR, PITA, and RNA22), scores were summed within genes or transcripts (if available) to calculate an aggregate score. For algorithms providing multiple transcript-level predictions (i.e., DIANA-microT-CDS, miRanda-MicroCosm, and TargetSpy), the transcript with the best score was selected as the representative transcript isoform. In all cases, predictions with gene symbol or RefSeq ID formats were translated into FlyBase format. To avoid testing and training our context model on the same data, we generated cross-validated predictions for the context model. To do so, we held out each transfection dataset, fit a linear regression model using the data from the remaining 5 datasets, and generated predictions on the held-out data.

### Microarray processing

We downloaded raw Affymetrix data measuring the effects of a miR-14 knockout (GEO accession GSE20202) (Varghese et al. 2010), a miR-34 knockout (day 20, GEO accession “GSE25008”) (Liu et al. 2012), and a miR-277 knockout (ArrayExpress accession “E-MEXP-3785”) (Esslinger et al. 2013) and processed data as previously described (Agarwal et al. 2015), with the exception that the *drosophila2FLYBASE* function in the “drosophila2.db” R Bioconductor package was used to map Affymetrix probe IDs to FlyBase IDs.

## DATA ACCESS

Raw RNA-seq and 3P-seq data were deposited in the NCBI Gene Expression Omnibus (GEO, accession number GSE74581). All associated scripts necessary to reproduce most of the figures of this paper are provided at: https://github.com/vagarwal87/TargetScanTools.

## ACKNOWLEDGMENTS

We thank Calvin Jan for contributing 3P-seq data for S2 cells, and other members of the Bartel lab for helpful discussions. This material is based upon work supported under a National Science Foundation Graduate Research Fellowship (to V.A.), a NIH Medical Scientist Training Program fellowship T32GM007753 (A.O.S.), an EMBO long-term fellowship (to I.U.), and NIH grants GM067031 and GM118135 (to D.P.B.). D.P.B. is an investigator of the Howard Hughes Medical Institute.

## AUTHOR CONTRIBUTIONS

V.A. carried out computational analyses and produced Github code, A.O.S. performed *Drosophila* transfections and associated experiments, P.T. implemented revisions to the TargetScanFly website, and I.U. annotated 3′-UTR isoforms using 3P-seq data. V.A., A.O.S., and D.P.B. conceived of the project, and V.A. and D.P.B. wrote the paper.

## SUPPLEMENTAL TABLES

**Table S1.** Processed mRNA abundances (measured in Fragments Per Kilobase Per Million, FPKM) and mRNA fold changes corresponding to each of the six miRNA transfection datasets.

**Table S2.** The 91 seed families broadly conserved in Drosophila species, listing for each family the miRNA names, seed sequences, and signal-to-background ratios for 5′-UTR and 3′-UTR sites. These ratios are plotted in Figure 2F. Families conserved since the ancestor of bilaterian animals are also indicated.

**Table S3.** Scaling parameters used to normalize data to the [0, 1] interval. Provided are the 5^th^ and 95^th^ percentile values for continuous features that were scaled, after the values of the feature were determined and transformed as indicated (Table 1).

| Feature | 8mer |  | 7mer-m8 |  | 7mer-A1 |  |
| --- | --- | --- | --- | --- | --- | --- |
|  | 5th % | 95th % | 5th % | 95th % | 5th % | 95th % |
| 3p_energy | -4.740 | 0.000 | -3.950 | 0.000 | -3.935 | 0.000 |
| Other_sites | 0.000 | 1.400 | 0.000 | 2.750 | 0.000 | 2.000 |
| Len_3UTR | 1.957 | 3.190 | 1.960 | 3.144 | 1.962 | 3.165 |
| Len_ORF | 2.782 | 3.666 | 2.740 | 3.706 | 2.717 | 3.742 |
| SA | -4.933 | -0.791 | -5.767 | -0.800 | -5.464 | -0.939 |
| P<sub>CT</sub> | 0.000 | 0.891 | 0.000 | 0.825 | 0.000 | 0.760 |

**Table S4.** Coefficients of the trained context model corresponding to each site type, as shown in Figure 3E. Using these coefficients and corresponding scaling factors (Table S3), context scores of predicted targets can be computed as described (Agarwal et al. 2015).

|  | 8mer | 7mer-m8 | 7mer-A1 |
| --- | --- | --- | --- |
| (Intercept) | -1.570 | -1.085 | -0.628 |
| 3p_energy | 0.212 | 0.311 | 0.174 |
| Other_sites | -0.419 | -0.529 | -0.089 |
| Len_3UTR | 1.006 | 0.978 | 0.439 |
| Len_ORF | 0.866 | 0.292 | 0.211 |
| SA | -0.448 | -0.286 | -0.253 |
| P<sub>CT</sub> | -0.371 | -0.302 | -0.263 |

## SUPPLEMENTAL FIGURE LEGENDS

**Figure S1.**
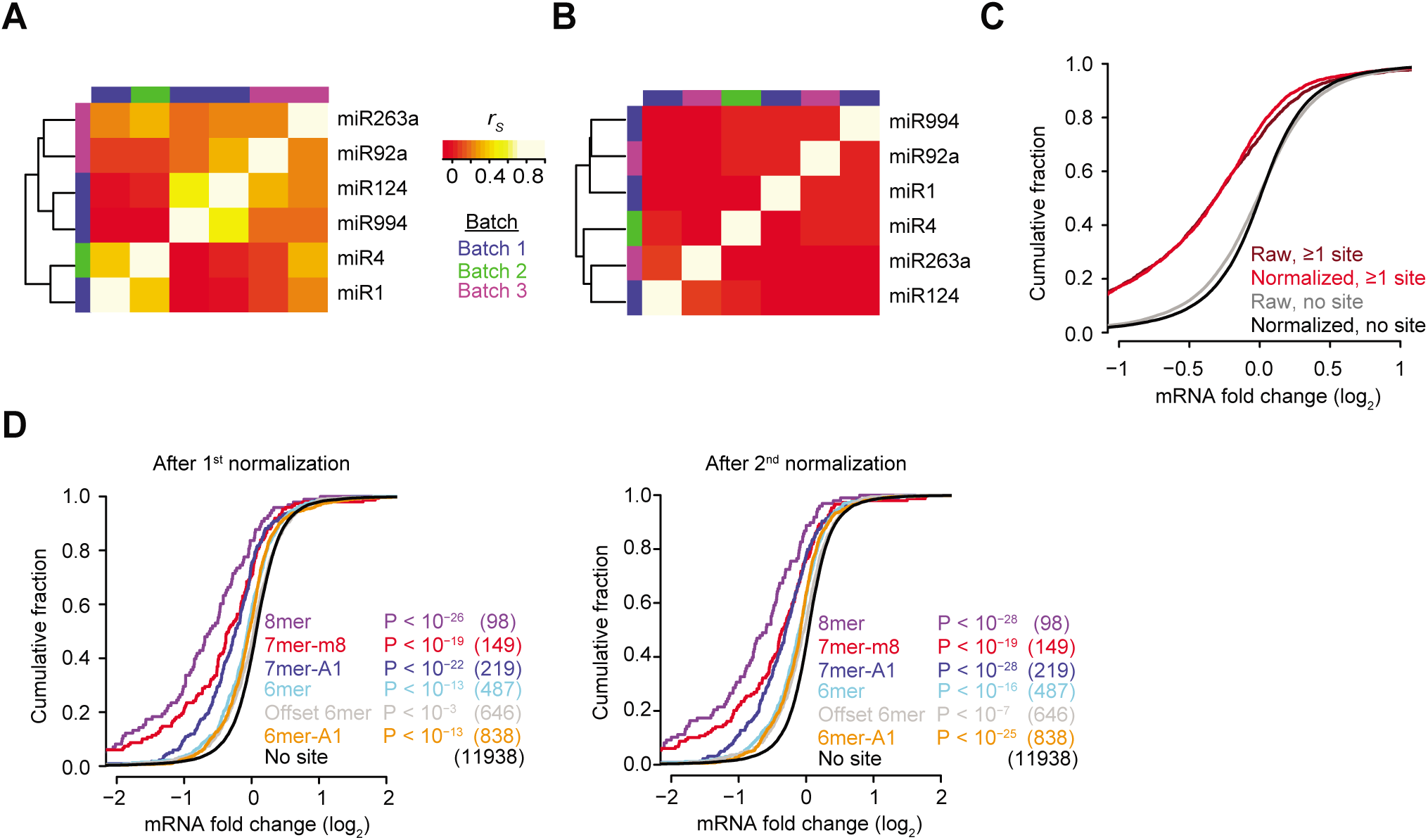
Preprocessing of RNA-seq datasets to minimize non-specific effects and technical biases. **(A)** Correlations observed between the responses of mRNAs without canonical 7–8-nt 3′-UTR sites to the transfected miRNAs. For each pair of experiments, the Spearman correlation (*r*_s_) of fold-change values for these mRNAs was calculated, and these *r*_s_ values, colored as indicated in the key, were then used for hierarchical clustering. The six transfection experiments were performed in three separate batches, which are colored as indicted in the batch list to show the correspondence between the clustering and the batches. **(B)** Reduced correlations observed between the responses of mRNAs without canonical 7–8-nt 3′-UTR sites to the transfected miRNAs after applying the PLSR technique. This heat map is as in (A) but plots the *r*s values obtained using PLSR-normalized mRNA fold changes. **(C)** Effects of the PLSR-based normalization on the fold-change distributions. Plotted are cumulative distributions of fold-changes observed after transfection of each of the six miRNAs, showing results for mRNAs containing either no site or at least one canonical 7–8-nt 3′-UTR site, either before (raw) or after PLSR-based normalization (normalized). **(D)** Residual mRNA fold changes either before (left) or after (right) a second round of normalization that removed biases between the mRNA fold changes and the A/U composition and sequence length of 5′ UTRs, ORFs, and 3′ UTRs. The panel showing results after the second round of normalization is the same as Figure 1C.

**Figure S2.**
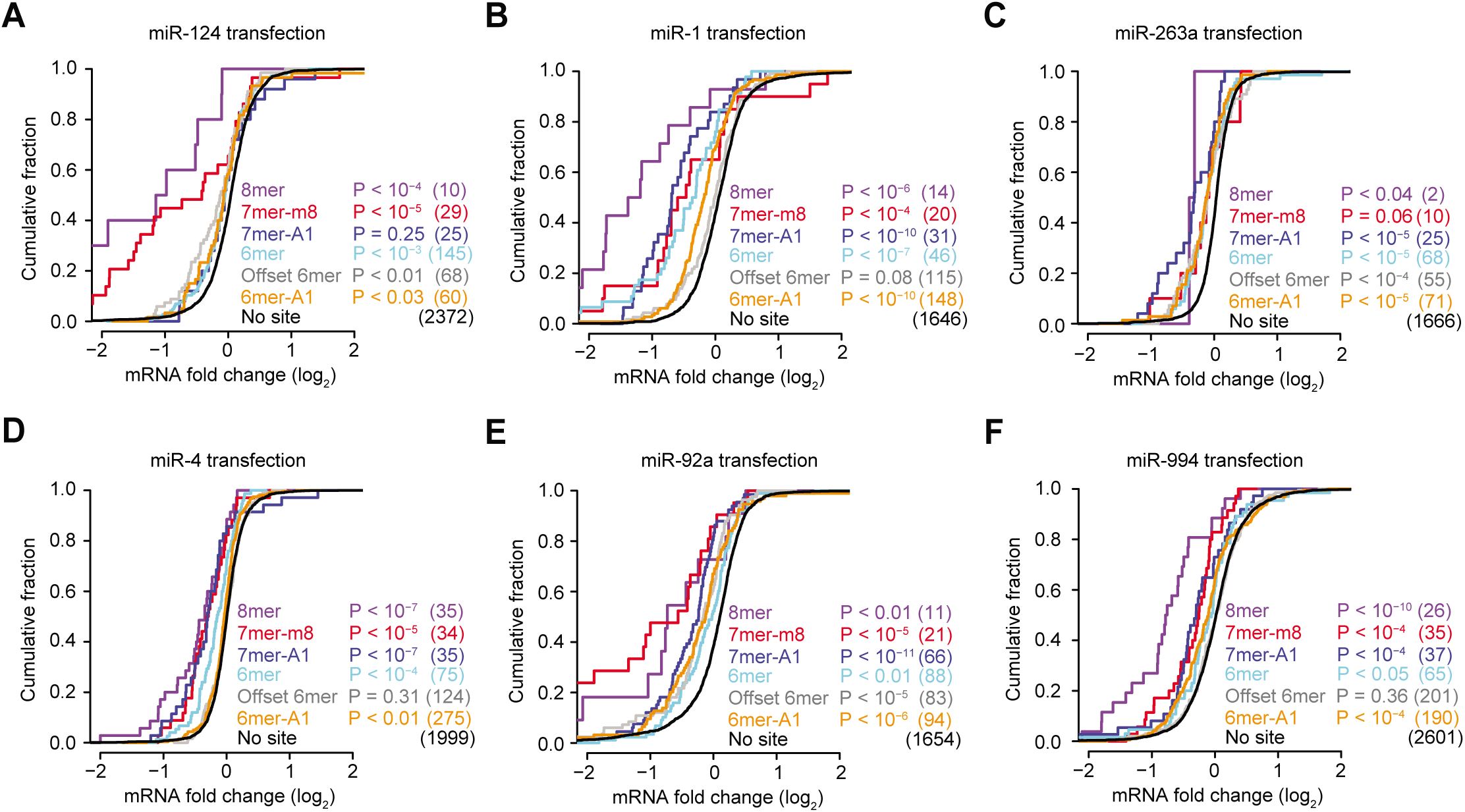
The efficacy of the canonical site types observed in Drosophila 3′ UTRs for individual experiments transfecting miR-124 **(A)**, miR-1 **(B)**, miR-263a **(C)**, miR-4 **(D)**, miR-92a **(E)**, or miR-994 **(F)**. Otherwise, these panels are as in Figure 1C.

**Figure S3.**
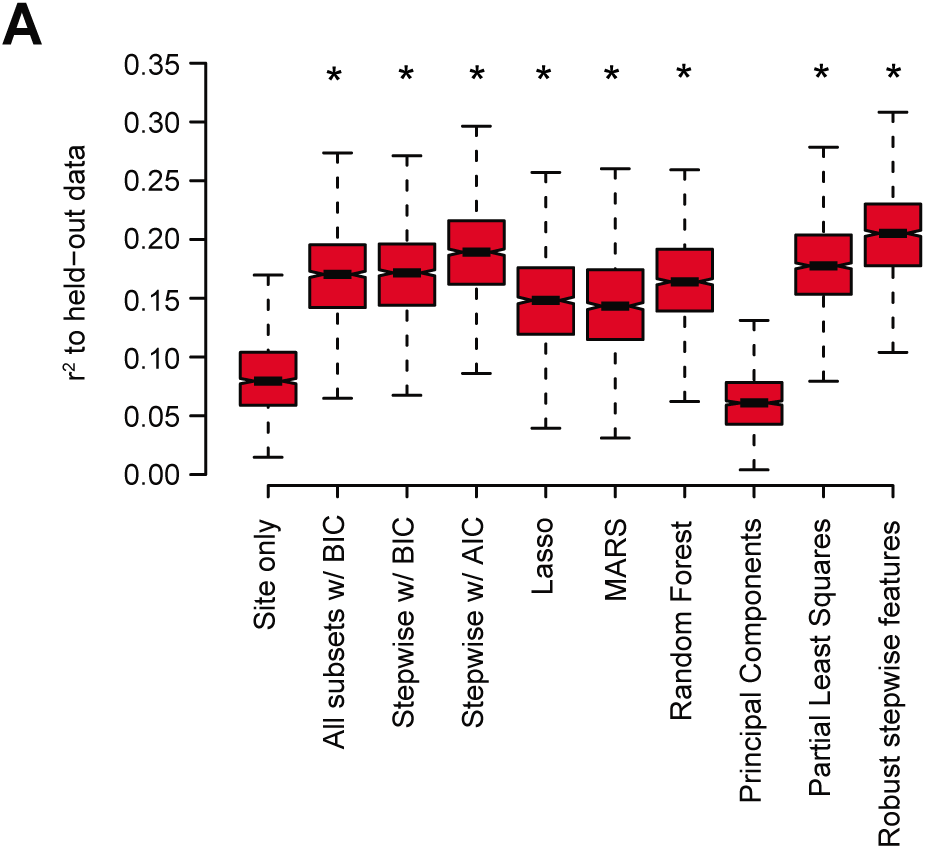
Performance of the models generated using different stepwise-regression methods compared to that of the site-only model. Shown are boxplots of r^*2*^values for each of the models across all 1000 sampled test sets. Highly significant improvement from the site-only model is indicated (\**P* < 10^−15^, paired Wilcoxon sign-rank test). Boxes indicate the median and interquartile ranges, and whiskers indicate either 1.5 times the interquartile range or the most extreme data point.

**Figure S4.**
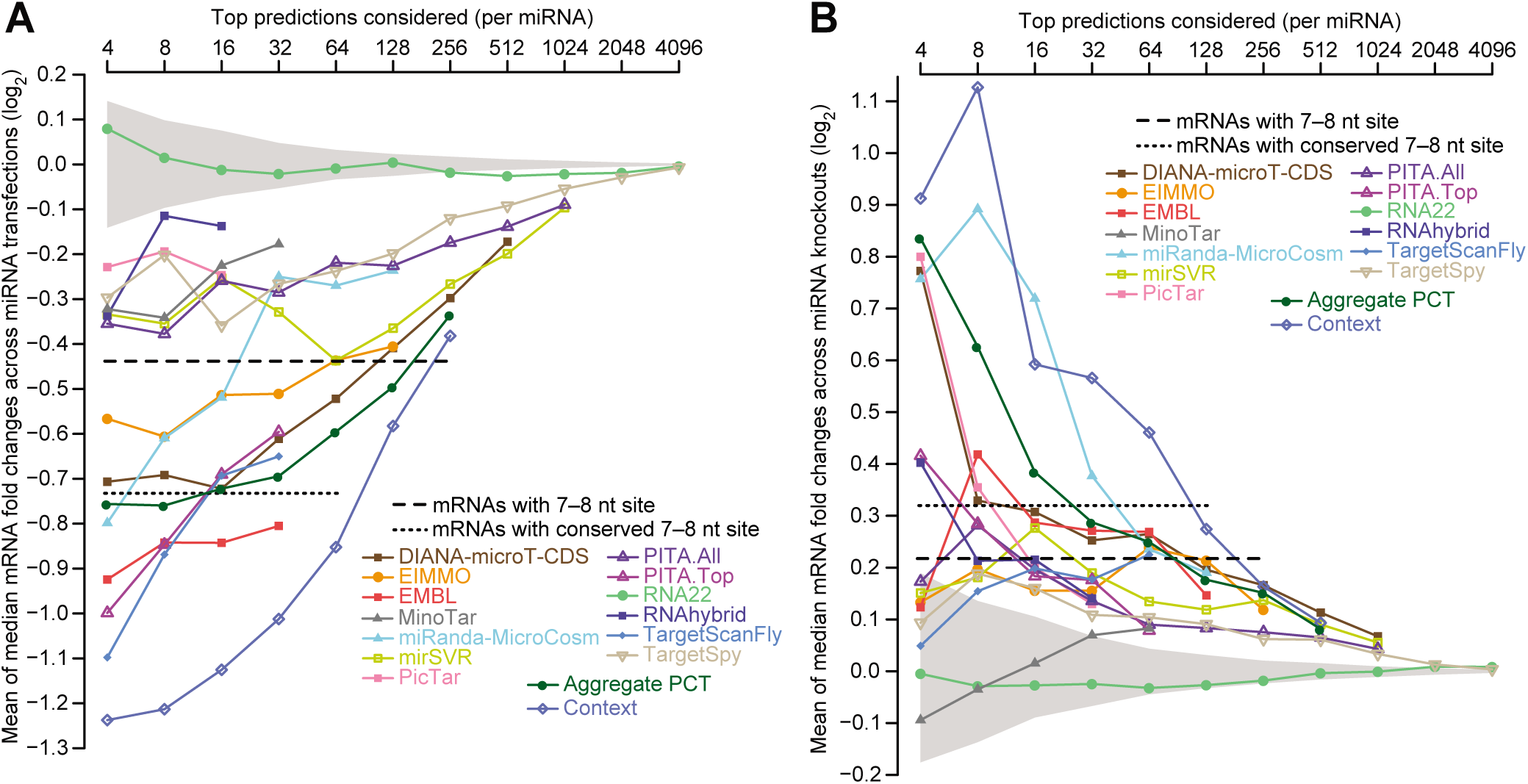
An alternative analysis of target-prediction performances in flies. **(A and B)** Evaluation of prediction performance plotting the mean of median values instead of the mean of mean values. Otherwise these panels are as in Figure 4A and B, respectively.

